# TranCEP: Predicting transmembrane transport proteins using composition, evolutionary, and positional information

**DOI:** 10.1101/293159

**Authors:** Munira Alballa, Faizah Aplop, Gregory Butler

## Abstract

Transporters mediate the movement of compounds across the membranes that separate the cell from its environment, and across inner membranes surrounding cellular compartments. It is estimated that one third of a proteome consists of membrane proteins, and many of these are transport proteins. Given the increase in the number of genomes being sequenced, there is a need for computation tools that predict the substrates which are transported by the transmembrane transport proteins.

In this paper, we present TranCEP, a predictor of the type of substrate transported by a transmembrane transport protein. TranCEP combines the traditional use of the amino acid composition of the protein, with evolutionary information captured in a multiple sequence alignment, and restriction to important positions of the alignment that play a role in determining specificity of the protein.

Our experimental results show that TranCEP significantly outperforms the state of the art. The results quantify the contribution made by each kind of information used.

## 1 Introduction

Transmembrane proteins are gates that organize a variety of vital cellular functions including cell signaling, trafficking, metabolism, and energy production. It is estimated that on average one in every three proteins in a cell is a membrane protein [Bue15, KST13]. Any defective or mis-regulated membrane protein can disturb the organism’s functioning, giving rise to disease [GO14]. About one-half of the drug targets today are membrane proteins such as transporters or related receptors [BRK17]. While the amino acid sequence of many membrane proteins is available, they are among the least characterized proteins in terms of their structure and function. For example, only 3% of the structures in the PDB are transmembrane proteins.

The publication of numerous genome projects has produced an abundance of protein sequences, many of which remain uncharacterized; transmembrane proteins are among the least characterized proteins, because experimental characterization of their structure and function is exceptionally difficult, owing to their hydrophobic surfaces and their lack of conformational stability. Consequently, there is an urgent need for computational approaches that use the available data to distinguish and characterize transmembrane proteins. Yet, this area of research is still in its early stages, and the researchers are far from finding a definitive solution.

Existing tools for the annotation of transporters that predict the substrate for the transport reaction lag behind tools for other kinds of proteins such as enzymes for metabolic reactions. Most tools predict the type of substrates [SCH10, COLG11, SH12, BH13, MCZ14], chosen from a small subset of substrate types, without attempting to predict the specific substrate, or predict the family or subfamily [GY08, LBUZ09, OCG10, BH13] for the protein within the Transporter Classification (TC) [BSJ03, SJTB06, SJRT^+^16], again without attempting to predict the specific substrate. For network modeling in systems biology [TP10, SAJT14], we require tools to process the complete proteome and predict each transport reaction; this means identifying the transport protein and the specific substrate.

Many tools rely simply on homology or orthology to predict transporters. This includes metabolic network tools merlin [DRFR15], and Pantograph [LZS15], as well as our system TransATH [AB17]. Amongst the tools for *de novo* prediction of substrate class, TrSSP [MCZ14] claims to be the state of the art.

Our previous efforts [Apl16] for *de novo* prediction of specific substrates for sugar transporters in fungi were not successful. However, from it we learned how much depends on very few residues of the transporter; often three or so residues, and often internal to different helix transmembrane segments (TMS) of the transporter [FBS^+^14]. These residues are far apart in the linear protein sequence, but are close to each in the three-dimensional structure of the protein when integrated in the membrane. In looking forward to how we might improve on approaches that rely on the amino acid composition of the protein, we developed a roadmap where the composition information would be combined with evolutionary information as captured by a multiple sequence alignment (MSA), and by positional information [TWNB12] about the residues responsible for determining specificity of the transporter. This roadmap is a schema for a large number of possible algorithms due to the many choices for encoding of amino acid composition, MSA algorithms, and algorithms for specificity determining sites [CC14]. We also realized the importance of the alignment preserving the TMS positions since the important residue positions seem to be located there. There are a number of such MSA algorithms [PFH08, CDTTN12, FTC^+^16, BGA^+^17].

Here, we conduct a preliminary study that shows the combination of information about protein composition, protein evolution, and the specificity determining positions has significant impact on our ability to predict the transported substrates. We chose the methodology and datasets of TrSSP [MCZ14] as our baseline, and varied this to illustrate the impact of each of the factors: compositional, evolutionary, and positional information. Our best approach, which defines the predictor we call TranCEP, for Transporter prediction using Compositional, Evolutionary, and Positional information, adopts the PAAC encoding scheme, the TM-Coffee MSA algorithm, and the TCS algorithm for determining informative positions in the MSA, to build a suite of Support Vector Machine (SVM) classifiers, one for distinguishing between each pair of classes of substrates.

### 2.2 Background

For most of the work done on the prediction of transport proteins [GO14], there is no available software, so it is difficult to reproduce the work and to compare the results of different articles.

*TransAAP* [RKP04] is a semi-automated analysis pipeline to input data into TransportDB. TransAAP targets only prokaryotes. A new genome is matched against the curated set of TransportDB proteins with assigned family using Blast with e-value cut-off of 1e-3. Information from these Blast searches against TransportDB are collected, as is information from searches against non-transporters in the nr protein database, and classification by orthology using COG. A web-based interface displays the information to help a human annotator decide and assign possible substrates or functions.

Pathway Tools includes the *Transport Inference Parser* (TIP) [LPK08] which analyses keywords in a gene annotation to assign Gene-Protein-Reaction associations to transport reactions in MetaCyc.

*G-Blast* [RS12] screens proteins against all entries in TCDB using Blast to retrieve the top hit, and HMMTOP to determine information about TMSs for the query and the hit sequence. It is an integral part of a manual protocol of Saier’s lab to predict the transport proteins for a genome [PVL^+^14].

The Zhao Lab has developed three methods: a nearest neighbour approach [LDZ08]; TransportTP [LBUZ09]; and TrSSP [MCZ14]. The nearest neighbour approach achieved a balanced accuracy of 67.0%.

*TransportTP* [LBUZ09] is a two-phase algorithm that combines homology and machine learning to predict TC family of one or more proteins. For training and cross-validation testing, *TransportTP* used the yeast proteome. For testing, it used 10 genomes from the TransportDB database [RCP07] of annotated prokaryote transporters. As an independent test, *TransportTP* is trained on the proteome of *A. thaliana* and then used to predict the transporters in four other plant proteomes. *TransportTP* achieved a balanced accuracy of 81.8%. Compared with the SVM-Prot classifier [LHC^+^06], on the five TC superfamilies and three families used by SVM-Prot, *TransportTP* achieved better recall and precision.

*TrSSP* (Transporter Substrate Specificity Prediction Server) [MCZ14] is a web server to predict membrane transport proteins and their substrate category. The substrate categories are: (1) oligopeptides (amino acid); (2) anion; (3) cation; (4) electron; (5) protein/mRNA; (6) sugar; and (7) other. *TrSSP* makes a top-level prediction of whether the protein is a transporter, or not. A SVM is applied with highest accuracy being reported using Amino Acid index (AAindex) and Position-Specific Scoring Matrix (PSSM).

The *disc function* system [GY08] uses amino acid composition and neural networks for discriminating channels/pores, electrochemical and active transporters, with an accuracy of 68%. When augmented with PSSM profiles and amino acid physicochemical properties they gained 5–10% in discrimination accuracy [OCG10]. They also considered six major families in TCDB [OCG10] with an average accuracy of 69%.

*TTRBF* [COLG11] considers four major classes of substrates: (i) electron, (ii) protein/mRNA, (iii) ion and (iv) others. It is an ensemble system combining amino acid composition, dipeptide composition, physicochemical properties, PSSM profiles and radial basis function (RBF) networks.

Schaadt et al. [SCH10] used amino acid composition, characteristics of amino acid residues and conservation to detect transporters based on different substrates, amino acids, oligopeptides, phosphates and hexoses and showed an accuracy of 75% to 90%. They classified to four substrate categories: amino acid, oligopeptide, phosphate, and hexose. The number of characterized transporters in *A. thaliana* for the four substrates numbered from 13 to 17. They constructed a vector for each protein using various types of amino acid composition, AAC, PAAC, PseAAC, PsePAAC, MSA-AAC, and used Euclidean distance from the query protein’s vector to the known vectors to rank the substrate categories. They found that AAC did not yield accurate results. However, PAAC performed as well as the more complicated PsePAAC and MSA-AAC, yielding accuracy over 90%.

Schaadt and Helms [SH12] compared the similarity of transporters in TCDB and annotated transporters in *A. thaliana* using amino acid composition and classified the proteins into three families. By distinguishing the amino acid composition of TMS and non-TMS regions, they could classify four different families with an accuracy of 80%.

Barghash and Helms [BH13] performed a comparison of three different approaches (homology, HMMER, MEME) for predicting substrate category and predicting TC family. They used four substrate categories, metal ions, phosphate, sugar, and amino acid; and 29 TC families with the most numerous examples. The datasets are from *E. coli*, *S. cerevisiae*, and *A. thaliana*, consisting of the 155, 158, 177, respectively, proteins that had both a substrate annotation and TC family annotation that are experimentally determined.

*Pantograph* [LZS15] was designed [Loi12] for metabolic pathway reconstruction of yeasts such as *Yarrowia lipolytica* [LDNS12] which accumulates lipids in the peroxisome component of the cell. It specifically models the cellular components and the transport across the membranes in a reference template, called the *scaffold model*. The Pantograph method relies on a database of profile HMMs for fungal protein families and their annotations that is maintained at G´enolevures in Bordeaux. The Pantograph algorithm first assigns Gene-Protein-Reaction associations, and then decides which compartments and reactions to include in the draft model based on these associations.

The scaffold model, which is the reference template for Pantograph, was manually curated to include 421 transport reactions. The associated transport protein families of orthologs were manually identified in the G´enolevures collection. The Pantograph software, written in Python, is available at http://pathtastic.gforge.inria.fr/. The distribution includes the scaffold model in SBML (Systems Biology Markup Language).

The *merlin* [DRFR15] system for the reconstruction of metabolic networks handles eukaryote genomes, and includes the determination of transport Gene-Protein-Reaction associations, as well as localization of reactions across a number of compartments: mitochondrion, endoplasmic reticulum (ER), and Golgi apparatus. In *merlin*, transport proteins are predicted based on the existence of TMS as predicted by TMHMM, and by similarity to entries in TCDB using the Smith-Waterman algorithm. The association of transport reactions and specific substrates for the predicted transport proteins is taken from a manually curated database of some 4000 TCDB entries originally. It now incorporates *TRIAGE* (Transport Reactions Annotation and Generation) [DGV+17] which contains information for 5,495 TCDB entries. The *merlin* software is available as open source Java code (http://www.merlin-sysbio.org).

*TransATH* [AB17] (Transporters via Annotation Transfer by Homology) is a system which automates Saier’s protocol based on sequence similarity. *TransATH* includes the computation of subcellular localization and improves the computation of transmembrane segments. The parameters of TransATH are chosen for optimal performance on a gold standard set of transporters and non-transporters from *S. cerevisiae*. A website http://transath.umt.edu.my for TransATH is available for use.

*SCMMTP* [LVY^+^15] uses a novel scoring card method (SCM) that utilizes the use of dipeptide composition to identify putative membrane transport proteins. The SCMMTP method first builds an initial matrix of 400 dipeptides and uses the difference between compositions of positives and negatives as an initial dipeptide scoring matrix. This matrix is then optimized using a genetic algorithm. *SCMMP* achieved an overall accuracy of 76.11% and Matthews correlation coefficient (MCC) of 0.47 in the independent dataset.

*Li et al* [LLX^+^16] used a SVM to predict substrate classes of transmembrane transport proteins by integrating features from PSSM, amino acid composition, biochemical properties and Gene Ontology (GO) terms. They achieved an overall accuracy of 80.45% on the independent dataset.

## 2 Materials and methods

### 2.1 Datasets

We used the same training dataset and test dataset as TrSSP [MCZ14] available at http://bioinfo.noble.org/TrSSP taken from Swiss-Prot (release 2013-03). Table 1 shows the statistics for the seven substrate classes considered: *amino acid*, *anion*, *cation*, *electron*, *protein/mRNA*, *sugar*, and *other*. The class *other* refers to transporters that do not belong to any of the other six classes. The dataset consisted of 760 transporters and 120 for the test dataset.

**Table 1.** The dataset.

| Substrate class | Training dataset | Test dataset |
| --- | --- | --- |
| Amino acid | 70 | 15 |
| Anion | 60 | 12 |
| Electron | 60 | 10 |
| Cation | 260 | 36 |
| Protein/mRNA | 70 | 15 |
| Sugar | 60 | 12 |
| Other | 200 | 20 |
| **Total** | 780 | 120 |

### 2.2 Databases

We use the Swiss-Prot database when searching for similar sequences. TM-Coffee uses the UniRef50-TM database, which consists of the entries in UniRef50 that have keyword *transmembrane*, when constructing multiple sequence alignments.

The dataset was derived from the Swiss-Prot database, so we make sure to remove all entries for the dataset from the two databases that we use.

### 2.3 Algorithm

Figure 1 ilustrates the steps of TranCEP. The sequence (a) in this case has four transmembrane segments (TMS) as shown by the gray shading. The example focusses on the first TMS, and abbreviates the middle section of the sequence. Part (b) shows an MSA conserving the TMS structure constructed by TM-Coffee, where the gray shading indicates the TMS location. Part (c) shows the colour coding of the reliability index of each column as determined by TCS, and shows how gaps replace unreliable columns in the filtered MSA. Part (d) shows a 400-dimensional vector of dipeptide frequencies (the PAAC composition) from the filtered MSA.

**Fig 1.**
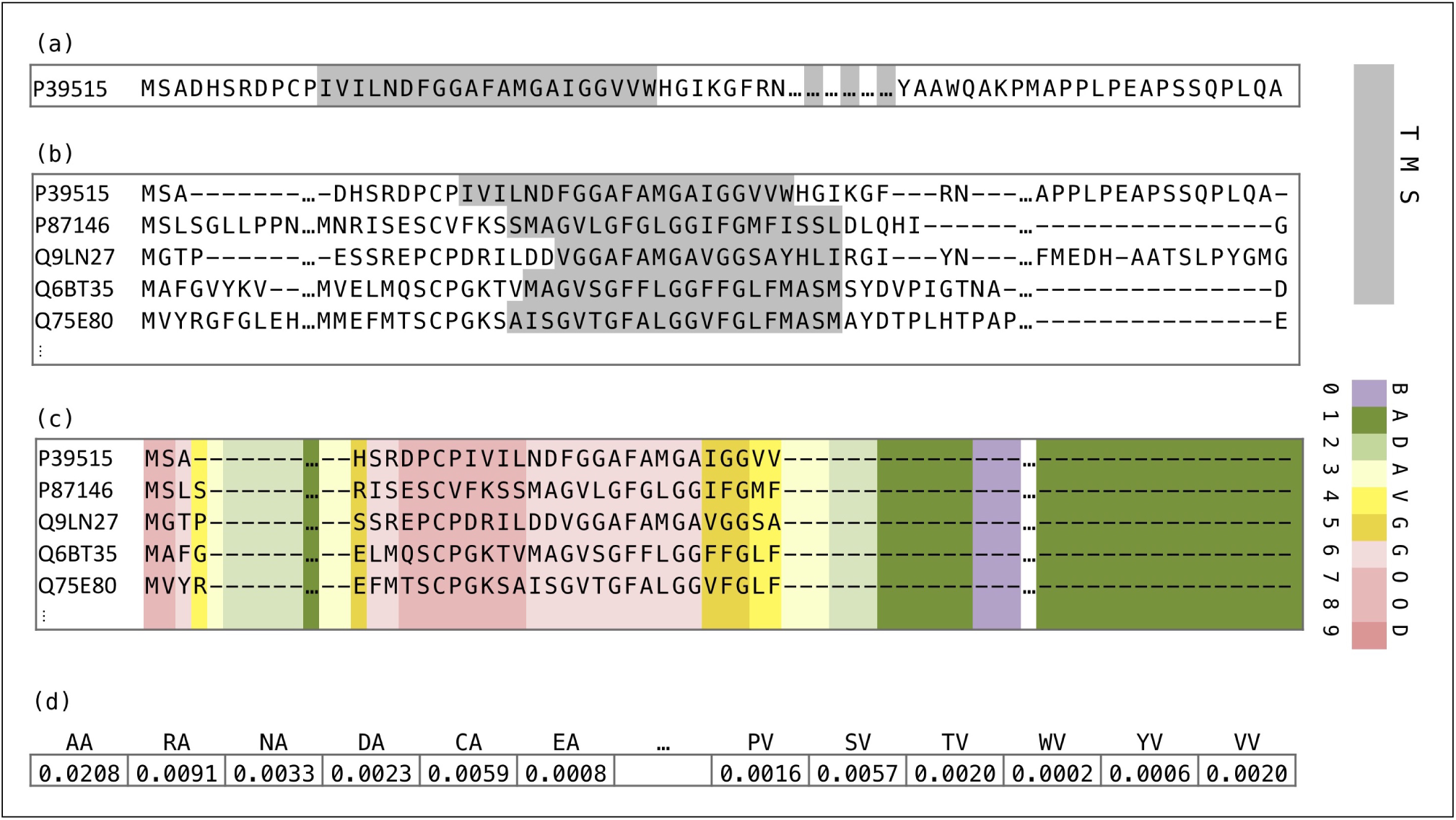
Example of Steps of TranCEP. The ilustrates the steps of TranCEP. Note that we abbreviate the middle section of the sequence. Part (a) shows the sequence the four TMS in gray shading. Part (b) shows an MSA constructed by TM-Coffee. Gray shading indicates a TMS. Part (c) shows the colour coding of the reliability index of each column as determined by TCS, and shows gaps in unreliable columns in the filtered MSA. Part (d) shows a 400-dimensional PAAC vector from the filtered MSA.

The template for combining evolutionary, positional, and composition information is presented in Algorithm 1. In this work, we use TM-Coffee to compute multiple sequence alignments (MSA) that conserve the transmembrane segments, and TCS to determine a reliability index for each position (column) in the MSA. We experimented with three composition schemes: amino acid composition (AAC), pairwise amino acid composition (PAAC), and pseudo-amino acid composition (PseAAC); and the optional use of TM-Coffee (TMC) and TCS.

#### Algorithm 1 Template of algorithm for composition vector. function comp vec(seq *s*)

// *Evolutionary (E) Step, optional*

*Construct MSA from s*

// *Positional (P) Step, optional*

Determine informative positions (columns) in MSA Filter uninformative posiitions from MSA

// *Composition (C) Step, mandatory*

**return** vector encoding composition of filtered MSA

**end function**

A template for the algorithm showing the role of evolutionary (E), positional (P), and composition (C) information. Note that the use of evolutionary (E) and positional (P) is optional; and that if positional (P) information is used then it requires evolutionary (E) information in the form of a multiple sequence alignment (MSA). Note that if Step (E) not done, then Step (C) encodes the sequence *s*. Note that if Step (E) is done but Step (P) MSA is not done, then Step (C) encodes the MSA.

Algorithm 2 shows the composition vectors being used to build a set of classifiers; SVM classifiers in this case. Algorithm 3 shows the prediction algorithm.

#### Algorithm 2 Build SVM classifiers.

**Require:** a training set *T* of sequences labelled with classes *C*_1_, …, C_n_

**Ensure:** a set of SVM’s *svm*(*i, j*) distinguishing class *C*_*i*_ and *C*_*j*_

**procedure** Build_SVMs(*T* : set of seq; *svm*: set of SVM)

for all seq *s* in *T* **do**

*v*(*s*) ← COMP VEC(*s*)

**end for**

**for all** pair (*C*_*i*_, *C*_*j*_) of classes **do**

*svm*(*i, j*) ← SVM.build({*v*(*s*) : *s ∈ T* ∩ (*C*_*i*_ ∪ *C*_*j*_)})

end for

end procedure

The algorithm to build a set of SVM classifiers to distinguish between each pair of classes in the training set. The actual construction of each SVM is done by an SVMM package’s build function.

#### Algorithm 3 Prediction.

**Require:** test sequence *s*

**Require:** a set of SVM’s *svm*(*i, j*) distinguishing classes *C*_*i*_ and *C*_*j*_

**Ensure:** result is predicted class *C*_*p*_

**function** PREDICT_CLASS(seq *s*)

*v ←* COMP_VEC(*s*)

**for all** pair (*C*_*i*_, *C*_*j*_) of classes **do**

*c*(*i, j*) ← *svm*(*i, j*) applied to *v*

**end for**

*p ←* mode of *c*(*i, j*) for all pairs (*i, j*)

**return** *C*_*p*_

**end function**

The prediction algorithm that applies each of the SVMs, and takes the class that is predicted most often by the set of SVMs.

### 2.4 Encoding amino acid composition

The properties of the amino acids at each position in the protein sequence can be encoded into vectors that summarize the overall composition of the protein. We implemented three approaches to encoding amino acid composition: *AAC*, *PAAC*, and *PseAAC*.

The *Amino Acid Composition* (AAC) is the normalized occurrence frequency of each amino acid. The fractions of all 20 natural amino acids are calculated as:

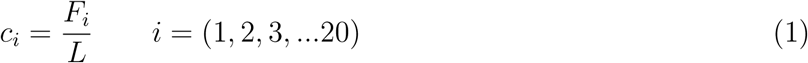

where *F*_*i*_ is the frequency of the *i*^*th*^ amino acid and *L* is the length of the sequence. The AAC is represented as a vector of size 20:

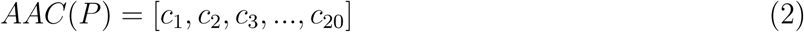

where *c*_*i*_ is the composition of *i*^*th*^ amino acid.

The *Pair Amino Acid Composition* (PAAC) is the normalized frequency of each pair of amino acids. The PAAC is calculated as

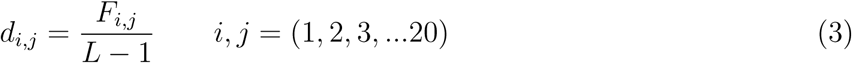

where *F*_*i,j*_ is the frequency of the *i*^*th*^ and *j*^*th*^ amino acids as a pair (dipeptide) and *L* is the length of the sequence. The PAAC is represented as a vector of size 400 as follows:

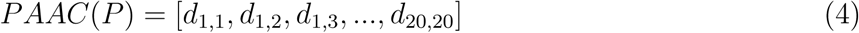

where *d*_*i,j*_ is the dipeptide composition of the *i*^*th*^ and *j*^*th*^ amino acid.

The *Pseudo-Amino Acid Composition* (PseAAC) [Cho01] combines the 20 components of AAC with a set of *sequence order correlation factors* that incorporates some biochemical properties. Given a protein sequence *R*_1_*R*_2_*R*_3_*R*_4_…*R*_*L*_ of length *L*, a set of descriptors called sequence order-correlated factors are defined as:

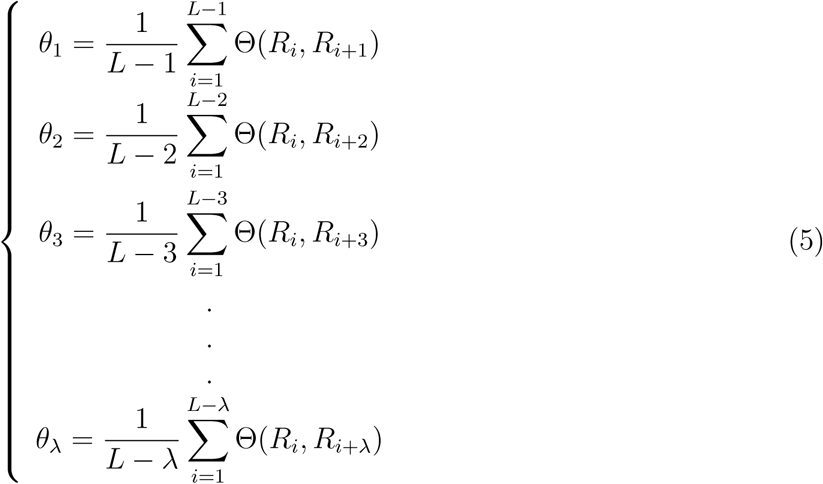

The parameter *λ* is chosen such that (*λ < L*). A correlation function is given by:

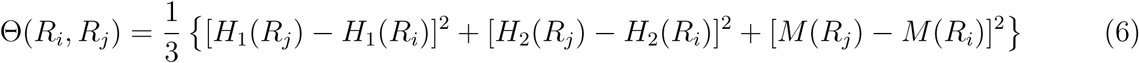

where *H*_1_(*R*) is the hydrophobicity value, *H*_2_(*R*) is hydrophilicity value, and *M* (*R*) is side chain mass of the amino acid *R*_*i*_. Those quantities are converted from their original values. For eaxmple, for hydrophobicity, *H*_1_(*R*) is derived from 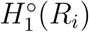, as follows:

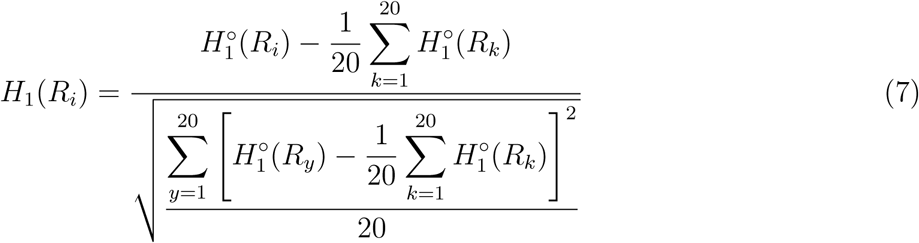

The original hydrophobicity value 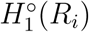 is taken from Tanford [Tan62]. The original hydrophilicity value 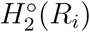 for the amino acid *R*_*i*_ is taken from Hopp and Woods [HW81]. The mass *M ^°^*(*R*_*i*_) of the *R*_*i*_ amino acid side chain can be obtained from any biochemistry text book.

PseAAC is represented as vector of size (20 + *λ*) as follows:

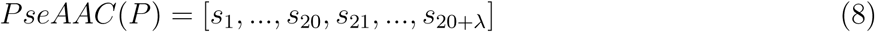

where *s*_*i*_ is the pseudo-amino acid composition

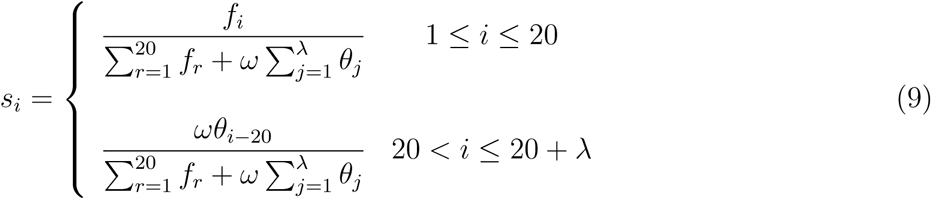

where *f*_*i*_ is the normalized occurrence frequency of the of the *ith* amino acid in the protein sequence, *θ*_*j*_ is the *j*^*th*^ sequence order-correlated factor calculated from Equation 5, and *ω* is a weight factor for the sequence order effect. The weight factor *ω* puts weight on the additional PseAAC components with respect to the conventional AAC components. The user can select any value from 0.05 to 0.7 for the weight factor. The default value given by Chou [Cho01] is .05.

### 2.5 Multiple sequence alignment

We adopted the *MSA-AAC* approach [SCH10] that combines amino acid composition with the evolutionary information available from a multiple sequence alignment (MSA). This is done by first retrieving homologous sequences of each protein sequence in the dataset, then building a MSA for the corresponding protein, then finally taking the counts for computing the composition information using all the residues in the MSA.

In [SCH10], they only used the *AAC* encoding, while we also applied the approach to the *PAAC* and *PseAAC* encodings. Another difference is that we adopted TM-COFFEE [CDTTN12] (Version-11.00.8cbe486) to compute alignments, rather than use ClustalW [THG94] as done in [SCH10], as we feel it is important to align the transmembrane segments well.

Other differences include searching the Swiss-Prot [BBA^+^03] database and retrieving a maximum of 120 homologous sequences instead of searching the non-redundant database nr and retrieving 1000 sequences. We sought to make the computational time more manageable because TM-COFFEE algorithm requires more memory usage and execution time.

Furthermore, all exact hits of the test sequences were removed from the Swiss-Prot and UniRef50-TM databases to maintain independence between the MSA and the test data.

Our alignment command was the following: :

~~~
t_coffee mysequences. fasta − mode psicoffee \
− protein_db uniref50 − TM \
− template_file PSITM
~~~

where mysequences.fasta contains the 120 similar sequences retrieved by BLAST.

### 2.6 Positional information

In order to focus the information on those positions in the protein that determine specificity, we need a method to determine those positions, and then to filter our information. The information is filtered by setting the entries for all other positions to null, that is, the symbol ‘-’ so that it is ignored when gathering counts for the amino acid composition information.

For determining the positions, we applied the *transitive consistency score* (TCS) algorithm [CDTN14] to the alignment. The TCS is a scoring scheme that uses a consistency transformation to assign a reliability index to every pair of aligned residues, to each individual residue in the alignment, to each column, and to the overall alignment. This scoring scheme is shown to be highly informative with respect to structural accuracy prediction based on benchmarking databases. The reliably index ranges from 0 to 9, where 0 is extremely uncertain and 9 is very reliable. Columns with a reliability index of below 4 were removed using the following command:

~~~
t_coffee − infile myMSA. aln −evaluate \
− output tcs_column_filter4. fasta
~~~

where myMSA.aln is the MSA file, tcs column filter4.fasta is the filtered file in FASTA format.

### 2.7 Training

Following TrSSP [MCZ14], we use a multi-class SVM with RBF kernel as implemented by the R e1071 library version 1.6-8 using a one-against-one approach in which 21 = (7 × 6)*/*2 binary classifiers are trained. The predicted class is determined through a voting scheme where all the binary classifiers are applied and the class that gets highest number of votes is the result. We also optimized both cost and gamma parameters of RBF kernel by performing grid search using *tune* function in the library (range of cost: 2 - 16, gamma: 2-e-5 - 1).

### 2.8 Methods

We implemented three approaches to encoding amino acid composition: *AAC*, *PAAC* as done by TrSSP [MCZ14], and *PseAAC*. These were followed by training using SVM to form the prediction methods **AAC**, **PAAC**, and **PseAAC** respectively.

By combining the amino acid composition and the evolutionary information obtained using TM-Coffee, followed by SVM, we implemented the prediction methods: **TMC-AAC**, **TMC-PAAC**, and **TMC-PseAAC** respectively.

Filtering was incorporated by applying TCS after TM-Coffee, then computing the amino acid composition vectors, and applying SVM. to implement the prediction methods: **TMC-TCS-AAC**, **TMC-TCS-PAAC**, and **TMC-TCS-PseAAC** respectively.

The method **TranCEP** is **TMC-TCS-PAAC**, as it gave the best performance during cross-validation.

### 2.9 Performance measurement

Four statistical measures were considered to measure the performance:

**sensitivity** which is the proportion of positives that are correctly identified:

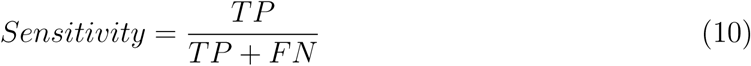

**specificity** which is the proportion of negatives that are correctly identified:

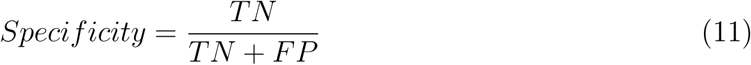

**accuracy** which is proportion of correct predictions made amongst all the predictions:

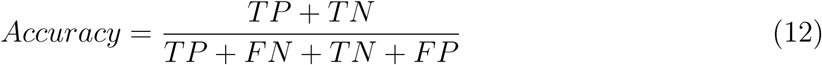

**Matthews correlation coefficient (MCC)** which is a single measure taking into account true and false positives and negatives:

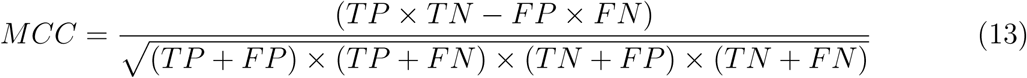

where *TP* is the number of true positives, *TN* is the number of true negatives, *FP* is the number of false positives, and *FN* is the number of false negatives.

We use the MCC because it is less influenced by imbalanced data and is arguably the best single assessment metric in this case [Din11, WP03, BDA13]. The MCC value ranges from 1 to −1, where 1 indicates a perfect prediction; 0 represent no better than random; and −1 implies total disagreement between prediction and observation. A high MCC value means the predictor has high accuracy on both positive and negative classes, and also low misprediction on both.

In the multi-class case with *K* classes, the MCC is calculated in terms of the confusion matrix *C* of dimension *K* × *K* [Gor04]:

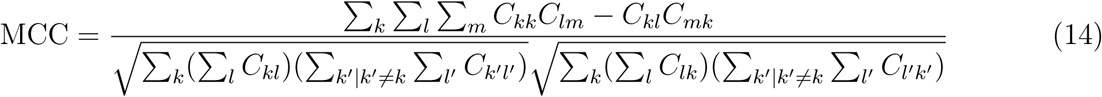

### 2.10 Experiments

Performance of each method was determined using both a five-fold cross-validation where the training dataset is randomly partitioned into five equal sized sets. A single set is kept as the validation data and the remaining four sets are used to train the SVM model. This model is then tested using the validation set. The cross-validation process is repeated four times where each of the sets is used once as the validation data. The performance of each model is averaged to produce a single estimation and the variation of performances is captured by computing the stranded deviation.

The performance of TranCEP on the test dataset was compared to the performance of TrSSP as reported in their paper [MCZ14].

## 3 Results and discussion

### 3.1 Cross-Validation

The performance of the nine methods were evaluated using five-fold cross-validation. Table 2 presents the overall accuracy and MCC of SVM models for the nine methods, sorted from the best to worst MCC. All SVM models that utilized evolutionary data notably performed better than the SVM models that did not. In addition, the top two models, **TMC-TCS-PAAC** and **TMC-TCS-AAC**, integrated positional and evolutionary information. The highest MCC was obtained by **TMC-TCS-PAAC**, which is the method chosen for our predictor **TranCEP**.

**Table 2.** Overall cross-validation performance of the methods. For each method, the table presents accuracy and MCC as *m±d*, where *m* is the mean and *d* is the standard deviation across the five runs of the cross validation.

| Method | Accuracy | MCC |
| --- | --- | --- |
| TMC-TCS-PAAC | 69.23±0.061 | 0.63±0.122 |
| TMC-TCS-AAC | 65.13±0.032 | 0.58±0.125 |
| TMC-PAAC | 63.33±0.044 | 0.51±0.129 |
| TMC-AAC | 63.08±0.036 | 0.50±0.092 |
| TMC-TCS-PseAAC | 63.30±0.055 | 0.49±0.129 |
| TMC-PseAAC | 61.79±0.029 | 0.49±0.131 |
| PseAAC | 47.69±0.024 | 0.27±0.104 |
| PAAC | 45.89±0.031 | 0.25±0.101 |
| AAC | 45.38±0.033 | 0.22±0.099 |

The performances of different SVM models were evaluated using five-fold cross-validation. The highest MCC was obtained by **TMC-TCS-PAAC** which is the method chosen for our predictor **TranCEP**. This method incorporates the use of PAAC with evolutionary data in the form of MSA with positional information, in which columns that have a reliability below 4 are filtered out. We found that the performance peaked using this threshold and started to decline when filtering columns with a higher reliability index. The **TMC-TCS-AAC** method yielded an overall MCC of .63. The detailed performance is presented in Table 3, MCC was 0.82, 0.61, 0.62, 0.58, 0.46, 0.52, and 0.69 for *amino acid*, *anion*, *cation*, *electron*, *protein/mRNA*, *sugar*, and *other*, respectively.

**Table 3.** Cross validation performance for TMC-TCS-PAAC. The table presents cross validation results for TranCEP, which is TCS-TMC-PAAC. For each substrate class, the table presents specificity, sensitivity, accuracy and MCC as *m±d*, where *m* is the mean and *d* is the standard deviation across the five runs of the cross validation.

| Class | Specificity | Sensitivity | Accuracy | MCC |
| --- | --- | --- | --- | --- |
| Amino acid | 98.82±0.004 | 80.84±0.054 | 96.06±0.017 | 0.82±0.042 |
| Anion | 99.30±0.007 | 47.89±0.199 | 93.33±0.014 | 0.61±0.119 |
| Cation | 86.66±0.022 | 79.70±0.043 | 81.50±0.039 | 0.62±0.081 |
| Electron | 96.58±0.030 | 63.13±0.118 | 92.01±0.019 | 0.58±0.098 |
| Protein | 97.21±0.018 | 52.00±0.087 | 90.93±0.031 | 0.46±0.036 |
| Sugar | 98.19±0.019 | 70.51±0.105 | 94.39±0.021 | 0.52±0.192 |
| Other | 83.02±0.035 | 67.23±0.198 | 76.43±0.068 | 0.69±0.178 |
| **Overall** |  |  | 69.23±0.061 | 0.63±0.122 |

Table 4 presents the confusion matrix for TranCEPwhen run on the independent test set. Most of the confusion occurs between a substrate class and the class *other*.

**Table 4.** Confusion matrix for TranCEP. The table presents the number of proteins in an actual substrate class that are predicted by TranCEP to fall in each substrate class. The independent test set is used.

| **Actual\Predicted** | Amino acid | Anion | Cation | Electron | Protein | Sugar | Other | **Total** |
| --- | --- | --- | --- | --- | --- | --- | --- | --- |
| Amino acid | 9 | 0 | 2 | 0 | 0 | 0 | 4 | 15 |
| Anion | 1 | 7 | 1 | 0 | 0 | 0 | 3 | 12 |
| Cation | 0 | 0 | 34 | 0 | 1 | 0 | 1 | 36 |
| Electron | 0 | 0 | 1 | 8 | 0 | 0 | 1 | 10 |
| Protein | 0 | 0 | 3 | 0 | 10 | 0 | 2 | 15 |
| Sugar | 0 | 1 | 0 | 0 | 0 | 8 | 3 | 12 |
| Other | 1 | 3 | 2 | 0 | 0 | 1 | 13 | 20 |

### 3.2 Comparison

Table 5 compares the performance of TranCEP to TrSSP [MCZ14] on the independent test set. For all substrate classes, TranCEP scored higher in accuracy, specificity, and MCC. However, for the true positive rate as measured by sensitivity, TrSSP performed better on the *cation* and *other* classes, and TranCEP either matched or outperformed TrSSP on the other five classes. Overall, TranCEP obtained MCC of 0.69 which is 68% higher than that of TrSSP.

**Table 5.** Comparing TranCEP and TrSSP. The table presents our data for TranCEP built with the training set and run on the test set. The corresponding results for TrSSP are taken from their original paper. The table shows specificity, sensitivity, accuracy and MCC for each of the seven substrate types; and the average accuracy and MCC. We calculated accuracy as proportion of correct predictions divided by the total number of predictions, and MCC from the confusion matrix as in Equation 14. The TrSSP results were calculated as the average across the seven classes; if we adopt the same method the average accuracy is 91.27% and the average MCC is 0.69.

| **Class** | **Specificity** |  | **Sensitivity** |  | **Accuracy** |  | **MCC** |  |
| --- | --- | --- | --- | --- | --- | --- | --- | --- |
|  | TrSSP | TranCEP | TrSSP | TranCEP | TrSSP | TranCEP | TrSSP | TranCEP |
| Amino acid | 82.42 | 98.10 | 93.33 | 60.00 | 83.33 | 91.75 | 0.49 | **0.66** |
| Anion | 69.05 | 96.30 | 75.00 | 58.33 | 69.44 | 90.82 | 0.23 | **0.56** |
| Cation | 74.31 | 89.29 | 75.00 | 94.44 | 74.44 | 89.00 | 0.41 | **0.78** |
| Electron | 91.78 | 99.05 | 80.00 | 80.00 | 91.11 | 97.80 | 0.50 | **0.88** |
| Protein | 82.42 | 99.07 | 93.33 | 66.67 | 83.33 | 93.68 | 0.49 | **0.75** |
| Sugar | 76.79 | 99.07 | 91.67 | 66.67 | 77.78 | 94.68 | 0.38 | **0.74** |
| Other | 73.13 | 86.00 | 60.00 | 65.00 | 71.67 | 80.91 | 0.23 | **0.44** |
| **Overall** |  |  |  |  | 78.88 | 74.17 | 0.41 | **0.69** |

### 3.3 Impact of Factors

Table 6 presents the impact of each factor — evolutionary information and positional information – on the overall MCC. The use of evolutionary information in form of MSA on the composition encodings **AAC**, **PAAC**, and **PseAAC** has improved the MCC by an average of 104% with the highest improvement being on **AAC** by 127%. The further use of positional information by filtering out the unreliable columns from the MSA has boosted the MCC of the composition encodings by an average of 132%. The impact of positional information over that already achieved by evolutionary information was an average of 14%. The highest impact was on **TMC-PAAC**, where the MCC improved by 24% in **TMC-TCS-PAAC**.

**Table 6.** Impact of factors on performance. The table notes the difference in MCC and the percentage improvement in MCC of the cross-validation performance for the introduction of evolutionary information using TM-Coffee, and positional information using TCS.

| Encoding<br>X | MCC |  |  | TMC-X to X |  | TMC-TCS-X to X |  | TMC-TCS-X to TMC-X |  |
| --- | --- | --- | --- | --- | --- | --- | --- | --- | --- |
|  | X | TMC-X | TMC-TCS-X | Delta | Percent | Delta | Percent | Delta | Percent |
| AAC | 0.22 | 0.50 | 0.58 | 0.28 | 127% | 0.36 | 164% | 0.08 | 16% |
| PAAC | 0.25 | 0.51 | 0.63 | 0.26 | 104% | 0.38 | 152% | 0.12 | 24% |
| PseAAC | 0.27 | 0.49 | 0.49 | 0.22 | 81% | 0.22 | 81% | 0.00 | 0% |
| **Average** |  |  |  | 0.25 | 104% | 0.32 | 132% | 0.07 | 13% |

Table 7 presents a breakdown on where the informative positions, as determined by TCS, are located with respect to the tramsmembrane segments (TMS). The protein sequence positions were divided into those in the TMS and those not in the TMS. Those in the TMS were divided into the interior one-third positions, and the remaining exterior positions in the TNS. The non-TMS positions were divided into those near a TMS, that is, within 10 positions), and the remaining positions far from a TMS. The location of the TMS was predicted using HMMTOP [TS01] to predict the *α*-helical TMS and PRED-TMBB2 [TEB16] for *β*-barrel TMS. Overall the positional information filtered out an average of (35% *±*7%) of the sequence. We found that *amino acid*, *anion*, *cation*, *sugar*, and *other*, substrates have significantly more important positions in the TMS compared with non-TMS and significantly more important positions close to TMS compared with positions far from TMS (Student’s t-test, P *<*0.0001). In contrast, in *protein/mRNA* and *electron* substrates, the difference was not significant. This could be because these two classes have fewer TMSs, and TCS did not capture their conservation.

**Table 7.**
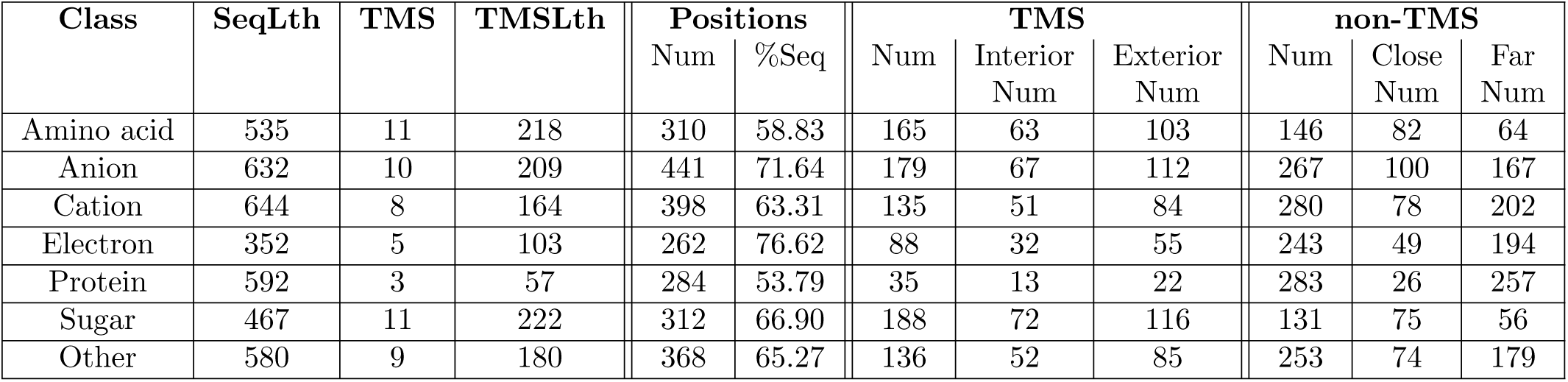
Positional information. The table present information on the sites retained by the TCS filtering step. For each class of substrates in the dataset, the table presents the average sequence length (**SeqLth**), the average number of TMS regions (**TMS**), and the average total number of residues in the TMS regions (**TMSLth**). It also presents the average of the number of positions retained by the filtering (**Positions:Num**), and that number as a percentage of the total sequence length (**Positions:%Seq**). It notes the total number of sites that occur in the TMS regions (**TMS:Num**), and the non-TMS regions (**non-TMS:Num**). For the TMS regions, it presents the average number of sites that occur in the interior central one-third of the TMS regions (**TMS:Interior:Num**), and in the remaining exterior regions outside the central one-third of the TMS regions (**TMS:Exterior:Num**). For the non-TMS regions, it presents the average number of sites that occur close to the TMS regions (within 10 positions of the TMS) (**non-TMS:Close:Num**), and the remaining sites far from the TMS regions (**non-TMS:Far:Num**).

| Class | SeqLth | TMS | TMSLth | Positions |  | TMS |  |  | non-TMS |  |  |
| --- | --- | --- | --- | --- | --- | --- | --- | --- | --- | --- | --- |
|  |  |  |  | Num | %Seq | Num | Interior<br>Num | Exterior<br>Num | Num | Close<br>Num | Far<br>Num |
| Amino acid | 535 | 11 | 218 | 310 | 58.83 | 165 | 63 | 103 | 146 | 82 | 64 |
| Anion | 632 | 10 | 209 | 441 | 71.64 | 179 | 67 | 112 | 267 | 100 | 167 |
| Cation | 644 | 8 | 164 | 398 | 63.31 | 135 | 51 | 84 | 280 | 78 | 202 |
| Electron | 352 | 5 | 103 | 262 | 76.62 | 88 | 32 | 55 | 243 | 49 | 194 |
| Protein | 592 | 3 | 57 | 284 | 53.79 | 35 | 13 | 22 | 283 | 26 | 257 |
| Sugar | 467 | 11 | 222 | 312 | 66.90 | 188 | 72 | 116 | 131 | 75 | 56 |
| Other | 580 | 9 | 180 | 368 | 65.27 | 136 | 52 | 85 | 253 | 74 | 179 |

## 4 Conclusion

We have developed a novel method TranCEP for *de novo* prediction of substrates for membrane transporter proteins that combines information based on amino acid composition, evolutionary information, and positional information. TranCEP incorporates first, the use of evolutionary information taking 120 similar sequences and constructing a multiple sequence alignment using TM-Coffee, second, the use of positional information by filtering to reliable positions as determined by TCS, and third, the use of pair amino acid composition. TranCEP achieved an overall MCC of 0.69, which is a 68% improvement over the state-of-the-art method TrSSP that uses the primary sequence alone. In addition, we evaluated the impact on performance of each factor: incorporating evolutionary information and filtering the unreliable positions. We observed that using amino acid composition alone does not obtain strong performance. The enhanced performance came from incorporating evolutionary and positional information. We learned that certain positions in the alignment have greater significance, and identifying them helped to boost the performance.

